# Cell state transition analysis identifies interventions that improve control of *M. tuberculosis* infection by susceptible macrophages

**DOI:** 10.1101/2023.02.09.527908

**Authors:** Shivraj M. Yabaji, Oleksii S. Rukhlenko, Sujoy Chatterjee, Bidisha Bhattacharya, Emily Wood, Marina Kasaikina, Boris Kholodenko, Alexander A. Gimelbrant, Igor Kramnik

**Affiliations:** The National Emerging Infectious Diseases Laboratories (NEIDL), Boston University, Belfield, Dublin 4, Ireland; Pulmonary Center, The Department of Medicine, Boston University School of Medicine. Belfield, Dublin 4, Ireland; Department of Microbiology, Boston University School of Medicine. Belfield, Dublin 4, Ireland; Systems Biology Ireland, School of Medicine and Medical Science, University College Dublin, Belfield, Dublin 4, Ireland; Conway Institute of Biomolecular & Biomedical Research, University College Dublin, Belfield, Dublin 4, Ireland; Department of Pharmacology, Yale University School of Medicine, New Haven, USA; Altius Institute for Biomedical Sciences, Seattle, WA, USA

**Author notes:** first co-authors (Shivraj M. Yabaji, Oleksii S. Rukhlenko).

## Abstract

Understanding cell state transitions and purposefully controlling them to improve therapies is a longstanding challenge in biological research and medicine. Here, we identify a transcriptional signature that distinguishes activated macrophages from TB-susceptible and TB-resistant mice. We then apply the cSTAR (cell State Transition Assessment and Regulation) approach to data from screening-by-RNA sequencing to identify chemical perturbations that shift the. transcriptional state of the TB-susceptible macrophages towards that of TB-resistant cells. Finally, we demonstrate that the compounds identified with this approach enhance resistance of the TB-susceptible mouse macrophages to virulent *M. tuberculosis*.

## Introduction

Understanding cell state transitions and purposefully controlling them to improve therapies is a longstanding challenge in biomedicine^1-6^. Immunity to infections requires that immunocompetent cells acquire and maintain cell states that optimally oppose specific patterns of microbial pathogenesis^7^. Successful pathogens, however, evolve mechanisms that allow them to induce cell states conducive to the pathogen’s survival and spread. The ability to identify distinct immune cell states and direct their transitions toward resistance-associated phenotypes is desirable for developing mechanistic immune therapies. Ideally, these interventions would take into consideration the individual host variation and stratify patients based on specific mechanisms of TB susceptibility of the genetic or environmental nature. To this end, recent studies used transcriptome analysis of peripheral blood and pathway analysis to identify aberrantly activated pathways in TB patients and stratified them according to mechanistic endotypes, suggesting the potential for endotype-specific therapies^5,6^.

An intracellular bacteria *Mycobacterium tuberculosis* (Mtb) co-evolved with humans for millennia and arguably is the most successful human bacterial pathogen: it has infected an estimated quarter of the human population, although tuberculosis disease (TB) is observed only in a minority of the infected humans^8-10^. In these, susceptible individuals, the bacterial infection leads to the formation of extensive lung pathology and pathogen transmission via aerosols^11-13^. Mechanisms underlying differences between the resistant and susceptible hosts are promising targets for immunomodulatory, so-called host-directed therapies^14,15^. Therefore, this is an area of intensive investigation^16^.

Macrophages play central roles in host resistance to virulent Mtb via effector and immunoregulatory mechanisms, including pathogen recognition, engulfment, and killing, as well as production of soluble mediators. The activation of appropriate effector mechanisms by macrophages is orchestrated by pathogen-derived ligands, cytokines produced by T lymphocytes, and other immune and stromal cells within the inflammatory milieu. Ideally, these signals induce activated macrophage states optimal for the pathogen elimination^17^. However, a number of the so-called intracellular pathogens survive inside macrophages and use them as a protected niche for the pathogen’s persistence and replication^18^.

Macrophage plasticity is exemplified by alternative activation states referred to as M1/M2 macrophage polarization. Canonically, these states are induced by cytokines: TNF and IFN*γ* induce the M1, and IL-4 or IL-13 induce the M2 macrophage polarization^19^. However, numerous observations provide evidence that this paradigm is insufficient to describe the entire palette of physiologically relevant macrophage states^20,21^. Although macrophages alternatively activated by IL-4 are, indeed, permissive for Mtb^22,23^, Mtb naturally induces Th1 T lymphocytes to produce M1 polarizing cytokines in both resistant and susceptible hosts in vivo, suggesting that M2 polarizing cytokines do not account for the differences in macrophage activation states associated with TB resistance and susceptibility^24,25^. Instead, hyperactivity of type I interferon (IFN-I) pathway in myeloid cells has been associated with TB susceptibility in humans and in animal models^26,27^. Whether the IFN-I hyperactivity is a biomarker of a specific macrophage state (polarization phenotype) remains unknown. Delineating pathways that shape and maintain specific macrophage phenotypes causally linked to TB susceptibility should enable rational design of interventions to correct them and avoid drugs that cause non-specific immune suppression^28,29^.

Previously we used forward genetic analysis to dissect mechanisms of TB progression in a mouse model of pulmonary TB and mapped a genetic locus, *sst1*, that specifically controlled the formation of human-like necrotic TB lesions in the lungs^30,31^. Subsequently, we found that the *sst1* locus-controlled macrophage activation by TNF, a cytokine essential for anti-tuberculosis immunity and granuloma formation^32^. During prolonged TNF stimulation, the *sst1*-susceptible macrophages develop escalating oxidative ^33^observed in their wild-type counterparts. Recently the *sst1*-encoded SP140 protein, which is not expressed in the *sst1*-susceptible macrophages, was shown to be the major determinant of the *sst1*-mediated susceptibility and type I interferon hyperactivity^34,35^, and its function was linked to maintaining heterochromatin silencing in activated macrophages^36,37^. This mechanism of action is consistent with our observation of coordinated change of multiple immune and metabolic pathways in the *sst1*-susceptible macrophages during persistent TNF stimulation^38^.

In this proof-of-principle study using the mouse model, we attempted to correct the *sst1*-susceptible macrophage phenotype underlying progression of pulmonary TB. First, we identified a differential gene expression signature distinguishing TNF response of TB-resistant (B6) and TB-susceptible (B6.Sst1S) macrophages. The 46 gene transcriptional signature was used for targeted mRNA expression analysis using bar-coded pooled RNA-seq for cost-efficient screening of candidate compounds in a medium-throughput format in 96-well plates. To determine the effects of specific pathway perturbations on the macrophage susceptibility state, these multi-dimensional data were analyzed using an explainable machine learning and dynamical analysis method, cell State Transition Assessment and Regulation (cSTAR)^33^. This approach has allowed us to identify specific states of macrophages in the transcriptomic space and quantify how drug perturbations affect the transitions between these states. Our findings suggested perturbations for switching from TB-susceptible to TB-resistant transcriptional states -such that the susceptible macrophages showed improved control of Mtb.

## Results

### I. Identification of a transcriptional signature of TNF response of the Mtb-permissive macrophages

Bone marrow derived macrophages (BMDMs) from naïve B6 (Mtb-resistant; R) and B6.Sst1S (Mtb-susceptible; S) mice were equally permissive for Mtb replication (**Fig.1A)**. In contrast, BMDM stimulated with TNF showed marked differences in Mtb resistance. TNF treatment had no effect on the bacterial control by the S macrophages, but significantly increased it in the R cells, as evidenced by reduced bacterial loads (**Fig.1A and B, Supplementary Fig.1A**). This agrees with our previous findings that an aberrant macrophage response to TNF underlies the *sst1*-mediated susceptibility to Mtb^32^. We posit that the functional differences in control of Mtb infection are reflected in distinct transcriptional states of TNF-stimulated S-and R-macrophages.

**Figure 1.**
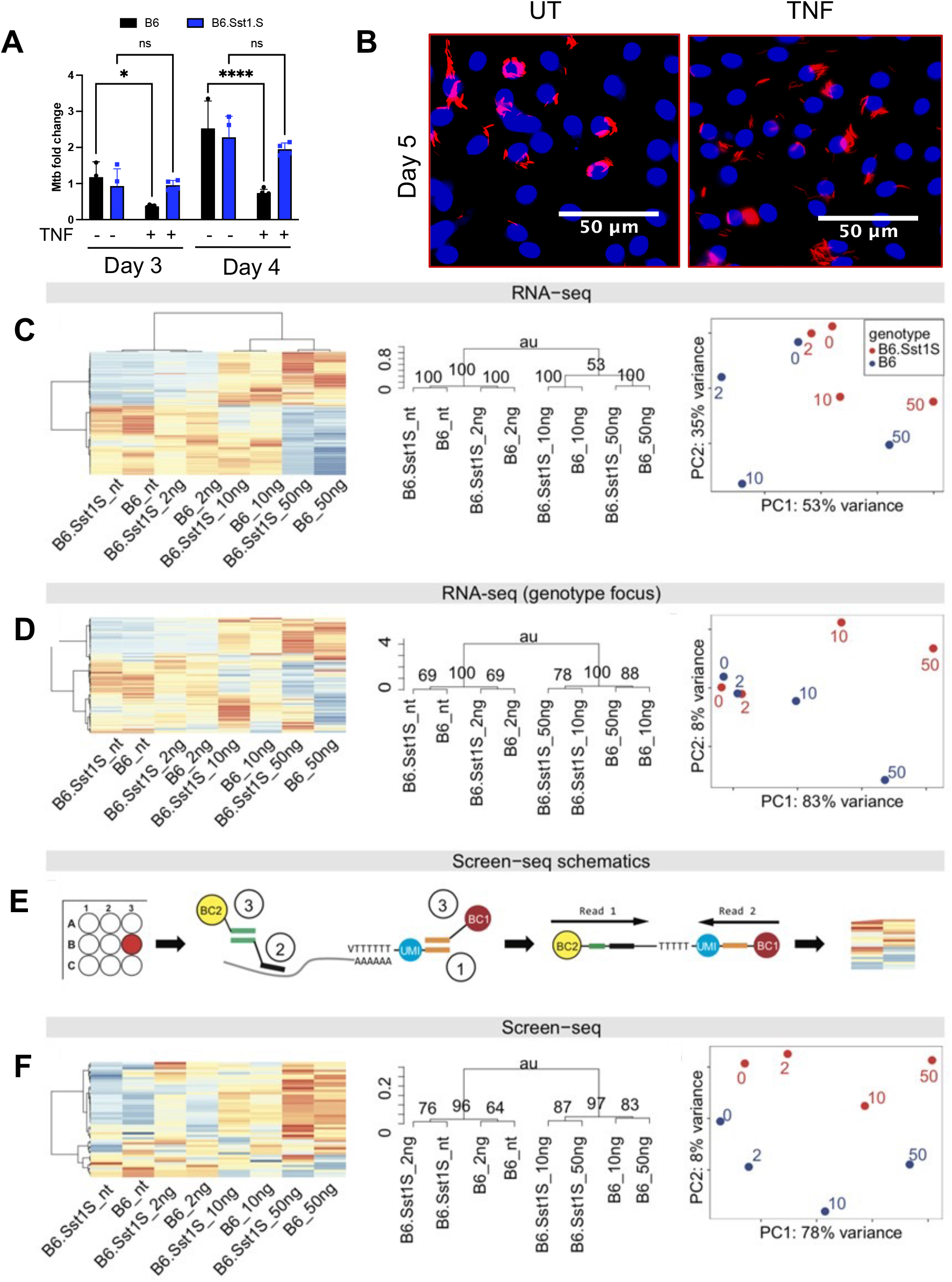
Transcriptional signature distinguishes the aberrant response of Mtb-susceptible macrophages. **A**. TNF improves control of Mtb in wildtype but not susceptible macrophages. B6 and B6.Sst1S BMDMs were treated with TNF (10 ng/ml) and subsequently infected with Mtb. The Mtb loads were determined using quantitative PCR (see Methods for details) on days 3 and 4 post infection. **B**. Fluorescent images of intracellular Erdman (SSB-GFP, smyc’::mCherry) in B6.Sst1S BMDMs treated with TNF compared to untreated control. All bacteria are marked in red (smyc’::mCherry) and nuclei are shown in blue (Hoechst 33342). BMDMs were infected with Erdman (SSB-GFP, smyc’::mCherry) at MOI = 1 and imaged at 5 days post infection. **C**. Whole-transcriptome macrophage response to TNF clusters by TNF dose. B6 and B6.Sst1S BMDMs were exposed to 0, 2, 10, and 50 ng/ml of TNF. Hierarchical clustering analysis of expression variation in bulk RNA-seq data: heatmap (left) and dendrogram (middle). P-values in the dendrogram computed by multiscale bootstrap resampling (“approximately unbiased”, P=100 is perfect confidence). Right panel - principal component analysis (PCA) of the same data. TNF concentration (in ng) is noted next to data points. Genetic background: *blue* – B6, *red* – B6.Sst1S. **D**. Genotype-specific response to TNF can be detected in RNA-seq data when using a subset of genes differentially expressed between B6 and B6.Sst1S BMDMs stimulated with 10 ng/ml TNF. Same data as in C but analyzed using only the top three hundred differentially-expressed genes. **E**. Overview of Screen-seq workflow (details in Methods). Left to right: Cells are lysed in-plate, and in each well, RNA is isolated using SPRI beads. Reverse transcription is performed using (1) oligo-dT primers with unique molecular identifiers (UMIs, *blue*) and common adapter (*orange*). Multiplex PCR is performed using (2) primers with gene-specific part (*black*) and common adapter (*green*); the reverse primer targets orange common adapter. Well-encoding is performed using (3) primers targeting common adapters coupled with barcodes (BC1 and BC2). Then all wells are pooled, Illumina sequencing adapters are added, and the pooled library is sequenced. In the profile analysis, Read 1 shows gene identity, UMI from Read 2 allows counting of cDNA molecules, while the barcodes BC1 and BC2 identify the original well. **F**. Screen-seq analysis distinguishes genotype-specific response to TNF. Same RNA samples as used for RNA-seq in C were subjected to Screen-seq with 46 targeted genes (see **Suppl. Table 1**). Same order of panels as in C and D.

To identify transcriptional signature of macrophage susceptibility, we compared whole transcriptomes of these macrophages using RNA-seq. Both R and S macrophages responded to TNF stimulation with large changes in gene expression (**Fig.1C**). When the whole transcriptome was analyzed, the effect of TNF far outweighed the effect of genotype, and the biological samples clustered by TNF concentration (**Fig.1C**). However, a subset of differences in transcriptional states of B6 and B6.Sst1S BMDMs could be found indicating that a targeted transcriptional profile can distinguish responses of S and R BMDMs to TNF (**Fig.1D**).

We then set out to identify perturbations that cause TNF-stimulated B6.Sst1S BMDMs to shift from the S to the R transcriptional state, aiming to use a targeted RNA sequencing approach, Screen-seq^39^, as outlined in **Fig.1E**. Briefly, cells are grown, treated and lysed in 96-well plates, with RNA isolated using magnetic beads. cDNA is synthesized using oligo-dT primers carrying unique molecular identifiers (UMI)^40^ and a common tail. Barcoded multiplexed amplicons from individual wells are pooled, and Illumina adaptors added for subsequent sequencing of the pooled library, with quantification of individual transcripts by UMI counting (see Methods for details).

For use in Screen-seq, we selected a set of genes induced by TNF, including (i) genes with differential expression between resistant and susceptible macrophages – both those with known roles in type I interferon response and proteotoxic stress response^32^, and those without such known roles, and (ii) a control set of genes responding to TNF but with no differential expression between R and S BMDMs. A 46 gene set of primers was designed to work in multiplex PCR (see Methods). When tested, they showed expression patterns similar to RNA-seq: UMI counts from Screen-seq analysis with these 46 genes recapitulated genotype-specific clustering of macrophage response to TNF stimulation (**Fig.1F**). We thus concluded that Screen-seq experimental and data-analysis pipeline could successfully resolve R and S expression profiles detectable in the transcriptome-wide data.

### II. Mapping transcriptional cell states in susceptible and resistant macrophages

To analyze the difference between TNF responses of R and S macrophages, we first applied cSTAR using the Screen-seq dataset of 46 gene responses^33^. In the transcriptomics dataspace we separated distinct macrophage states, using a classification machine learning method, the Support Vectors Machine (SVM). To visualize the results, we used Principal Component Analysis (PCA). **Fig. 2A** shows the projections of the screen-seq data points in a 46D space into a 2D space of the first two principal components (PC1 and PC2). The black line dividing the R and S macrophage states is a projection of the separating hyperplane in a 46D dataspace into a 2D PCA space. Reflecting their biology, the macrophage expression states are similar without TNF exposure (**Fig. 2A**).

**Figure 2.**
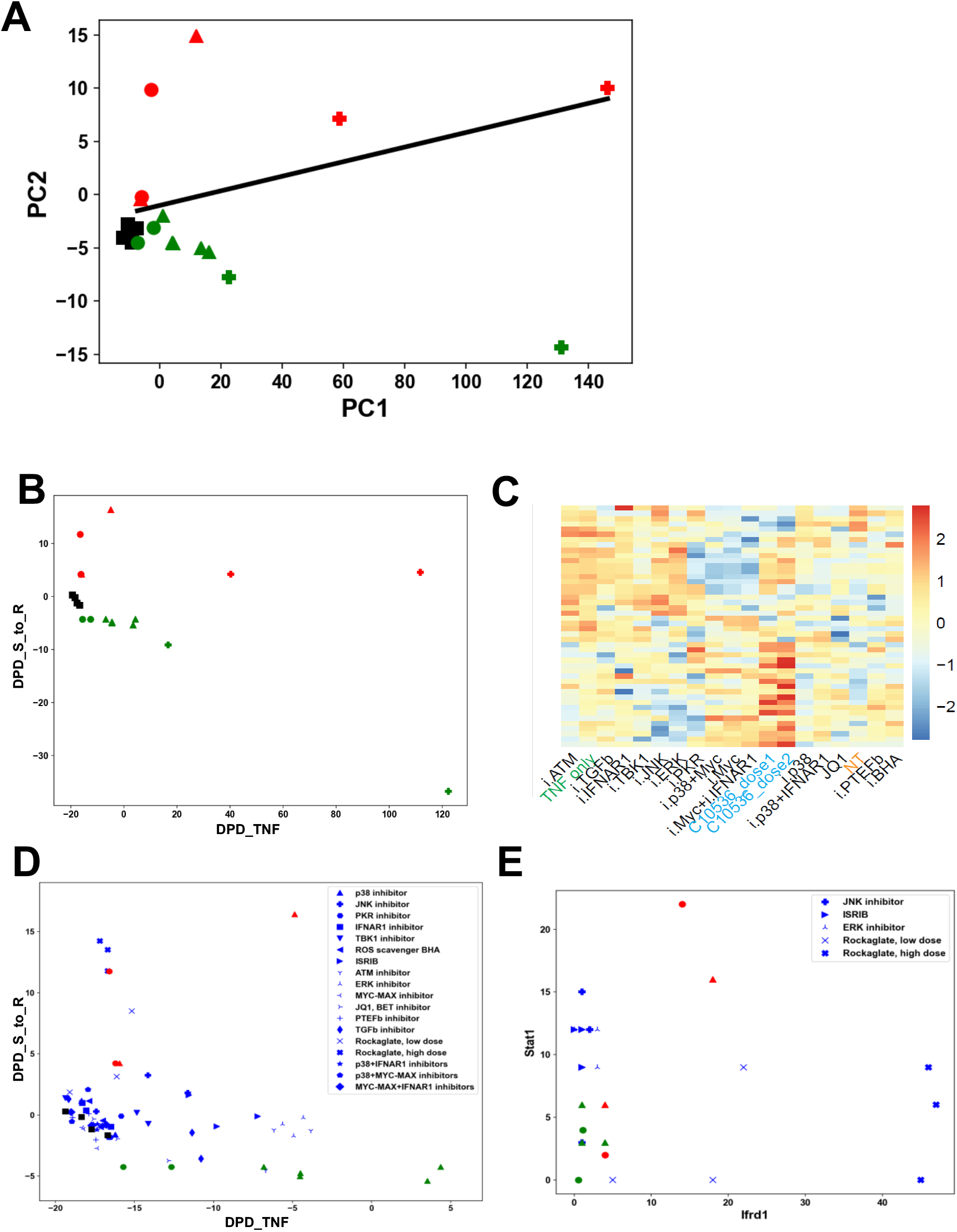
cSTAR analysis of targeted transcriptomics data. **A**. PCA representation of RNA-seq data from B6 (*red*) and B6.Sst1S susceptible mutant (*green*) BMDMs stimulated with different doses of TNF for 24 h: 2 ng/ml (*circles*), 10 ng/ml (*triangles*), 50 ng/ml (*pluses*), or unstimulated controls (*black squares). Black line* represents projection of SVM-generated maximum margin hyperplane to PCA-space. **B**. Same data as in A, represented in the Dynamic Phenotype Descriptors (DPDs) space. DPD_TNF quantifies the separation of TNF responses, DPD_TB quantifies the separation of resistant and susceptible phenotypes. Same colors and symbols as in A. **C**. Heatmap of targeted gene expression profiles (Screen-seq) of B6.Sst1S BMDMs after treatment with drugs for 24 h in the presence of 10 ng/ml TNF. Drugs listed on the X axis and in **Suppl. Table 2**. **D**. DPD space representation of the Screen-seq data. B6 (*red*) and B6.Sst1S (*green*) macrophages were treated with 2 ng/ml (*circles*) and 10 ng/ml (*triangles*) TNF for 24 hours, or untreated (*black squares*). *Blue* labels - B6.Sst1S BMDMs treated with 10 ng/ml TNF and selected drugs (**Suppl. Table 2**). **E**. Fold-changes of the *Stat1* and *Ifrd1 gene* expression in B6 (*red*) and B6.Sst1S BMDMs (*green*) treated with TNF 2 ng/ml (*circles*) or 10 ng/ml (*triangles*). *Blue* labels - B6.Sst1S BMDMs treated with 10 ng/ml TNF and selected compounds: rocaglate, ISRIB, JNK or ERK inhibitors, as denoted.

Although the PC visualization illustrates the SVM separation of the S and R macrophage states, it does not suggest ways to normalize transcriptomic patterns of the S states directing them towards the R states. Likewise, it cannot determine the direction of the changes in the transcriptomics patterns induced by the changes in the TNF doses. The cSTAR approach builds two STVs that direct desirable transitions between the macrophage states separated by the SVM (see Methods). The first STV indicates a path to move TB-susceptible to TB-resistant states (abbreviated as the STV_S_to_R_); the second STV shows the direction of transcriptomic changes between responses to low and high TNF doses (STV_TNF_). To understand how the difference in the genetic backgrounds and TNF doses affect macrophage states, we exploit two quantitative indicators of the corresponding phenotypic features: responses to TNF and TB susceptibility. These two phenotypic features are quantified by the Dynamic Phenotype Descriptors (DPDs) scores calculated as distances along the STV_TNF_ and the STV_S_to_R_, respectively (see Methods). Because the STV_S_to_R_ directs from S to R states, the DPD_S_to_R_ sign is positive at the R side of the separating border and negative on the S side. Likewise, the DPD_TNF_ sign becomes positive for high TNF doses. Instructively, the two STVs are nearly orthogonal in the transcriptomics space (the angle between the STV_TNF_ and the STV_S_to_R_ is about 103°) suggesting that molecular mechanisms of macrophage responses to TNF and TB susceptibility essentially do not overlap.

**In Fig. 2B** we visualized macrophage transcriptomic patterns for distinct genetic backgrounds and different TNF doses in the space of these two DPDs. This visualization suggests that TNF doses higher than 10 ng substantially decrease the DPD_S_to_R_ score for susceptible macrophages (cf. the data points for 10 and 50 ng) and weakly decrease it for resistant cells. At 10 ng TNF, the DPD_S_to_R_ score is maximal for resistant macrophages. These results suggest the optimal TNF dose of about 10 ng, which maximizes the DPD_S_to_R_ for resistant macrophages and does dramatically decreases the DPD_S_to_R_ for susceptible macrophages. Therefore, in subsequent experiments we were treating macrophages with 10 ng TNF.

### III. Use of Screen-seq dataset and cSTAR identifies perturbations normalizing the aberrant TNF response of the S macrophages

We applied perturbations to the S macrophages treated with 10 ng of TNFa and measured transcriptional responses of Screen-seq genes after 24 h (**Fig. 2C**). The small molecules for perturbations were selected based on our previous findings^29,32,38^ and included inhibitors of the IFN-I pathway (TBK1, IFNAR1 blocking antibodies), inhibitors of the MAP kinases (p38, JNK and ERK), PKR inhibitor, ROS scavenger BHA, Myc inhibitor and inhibitors of transcription elongation (pTEFb inhibitor and JQ1). We also used a selective inhibitor of cap-dependent protein translation, synthetic rocaglate CMLD010536. Full list of perturbations is given in **Supplementary Table 2**. Each perturbation was tested in three independent samples (wells).

#### Ranking drug perturbations by the STV and DPD

The perturbation outcomes were determined by the changes in the DPD_S_to_R_ and DPD_TNF_ scores. **Fig. 2D** visualizes these outcomes. **Supplementary Table 3** presents the DPD_S_to_R_ scores for S macrophages treated with 10 ng of TNF and different drugs and for R macrophages treated with 10 ng TNF (the DPD_S_to_R_ and DPD_TNF_ scores calculated for all perturbations can be found in **Supplementary Table 3**).

Each drug shifted TB-susceptible states towards TB-resistant states, judging by statistical significance, but only CMLD010536 treatments resulted in the crossing of the separating hyperplane (p≈0.003 for the CMLD010536 high dose), thus converting the S into R states **(Supplementary Table 3)**. The JNK and TBK inhibitors, a combination of p38 and Myc inhibitors, and integrated stress response inhibitor (ISRIB) also led to substantial phenotype movements, although neither of these treatments converted the S phenotype into R phenotype at the doses used.

Importantly, most of the applied drugs decreased the DPD_TNF_ scores, which shows that the TNF response of macrophages was partially suppressed by these drugs (**Fig. 2D**). In particular, a treatment with RocA moved the S transcriptomic patterns of macrophages stimulated with 10 ng TNF towards the R patterns as evidenced by the increasing DPD_S_to_R_ score, but it decreased the DPD_TNF_ scores to the values characteristic for 2 ng TNF. Treatments with ISRIB and a JNK inhibitor also increased the DPD_S_to_R_, whereas the concurrent DPD_TNF_ decrease was much lesser than for synthetic rocaglate CMLD010536. We hypothesize that if a drug combination increases the DPD_S_to_R_ score and simultaneously increases or, at least, does not strongly reduce the DPD_TNF_ score, it will synergistically improve transcriptomics state of susceptible macrophages.

To test this cojecture, we analyzed the outcome of drug perturbations in more detail. In the transcriptomics dataspace, each drug perturbation is determined by a perturbation vector, which connects the data points after and before the perturbation. The perturbation vector shows the fold-changes in each gene following a drug treatment. Considering the projection of the perturbation vector into the STV, we determine the contribution of each gene in the DPD change brought about by each drug. Ideally, a drug would activate the genes that are positive contributors to the STV_S_to_R_ and suppress the negative contributors, thereby normalizing the transcriptome. All three drugs, CMLD010536, JNK inhibitor (JNKi) and ISRIB, suppress *Ch25h, Fzd7, Hspa1b*, and *Sh2d3c*, which are negative STV_S_to_R_ contributors (**Supplementary Table 4**) and activate different genes with positive contributions to the STV_S_to_R_. CMLD010536 strongly activates *Ifrd1*, while JNKi and ISRIB activate *Stat1* and *Gsdmd*. This provides additional evidence for *potential* synergy between CMLD010536 and JNKi or ISRIB (**Fig. 2E**).

### IV. RNA-seq analysis of Mtb-infected B6.Sst1S macrophages treated with individual compounds and their combinations

Because TNF treatment is beneficial to counteract Mtb infection in resistant macrophages (**Fig.1A**), we hypothesized that a combination of rocaglate with ISRIB or JNKi should be advantageous for the transcriptome normalization in TNF-stimulated susceptible macrophages and, thus, improve their response to Mtb infection. To test our predictions, we performed an in-depth analysis of S-macrophages treated with suggested drug combinations and infected with virulent Mtb. In these experiments, we used a less toxic plant derived Rocaglamide A (RocA) to substitute for synthetic rocaglates used previously (**Supplementary Fig.1B, C and D**).

The S macrophages were stimulated with TNF and infected with Mtb, as above, and treated with RocA, ISRIB, SP600125 or their combination before and during the course of infection. We have chosen day 3 for the RNAseq measurements, since this is a critical point that determines the subsequent trajectory of the Mtb – macrophage interactions in our in vitro model (**Fig.1A**). Whole-cell transcriptome patterns of the R and S macrophages stimulated with TNF were separated using the entire RNA-seq data (accession number GSE115905). As described above, after building the STV_S_to_R_, we calculated the DPD_S_to_R_ score for the Mtb-infected S macrophages under all conditions. To get mechanistic insights, we determined each gene contribution to the change in the DPD_S_to_R_ score brought about by a single drug or a drug combination. These contributions are calculated as the projection of gene expression changes into the STV_S_to_R_ and quantify how each gene promotes or reverses a transition of S to R states (see Methods for details). We analyzed these data using two different approaches. First, we selected names of the genes that explain 80% of the change in the DPD_S_to_R_ score following a drug perturbation, and used Gene Ontology, KEGG, Reactome and STRING databases to annotate them. Second, for these genes we input the projections of their expression changes onto the STV_S_to_R_ together with their names into GSEA^41^ to obtain mechanistic insights into processes contributing to the TB resistance and susceptibiliuty. Below, we present the results of this analysis.

The changes in the DPD_S_to_R_ suggest that TNF pretreatment might have an overall positive effect on S macrophages infected with Mtb: it decreased the expression of many genes, which are negative contributors to the the DPD_S_to_R_ score, and, hence, their suppression increased this score and partially normalized the transcriptome of Mtb-infected macrophages (**Fig. 3A**). The GSEA analysis suggested that TNF changed the Interferon response to increase TB resistance, but TNF-induced changes in other genes contributed to the decrease of the DPD_S_to_R_, suppressing resistance The latter included several chemokines and chemokine receptors, genes related to cell cycle (cyclins A2 and B2) and DNA replication (**Fig. 3B**). Although the overall change in the DPD_S_to_R_ after TNF pretreatment showed a tendency towards resistance by the susceptible macrophages, this effect is insufficient to improve their ability to control Mtb (**Fig. 1A**).

**Figure 3.**
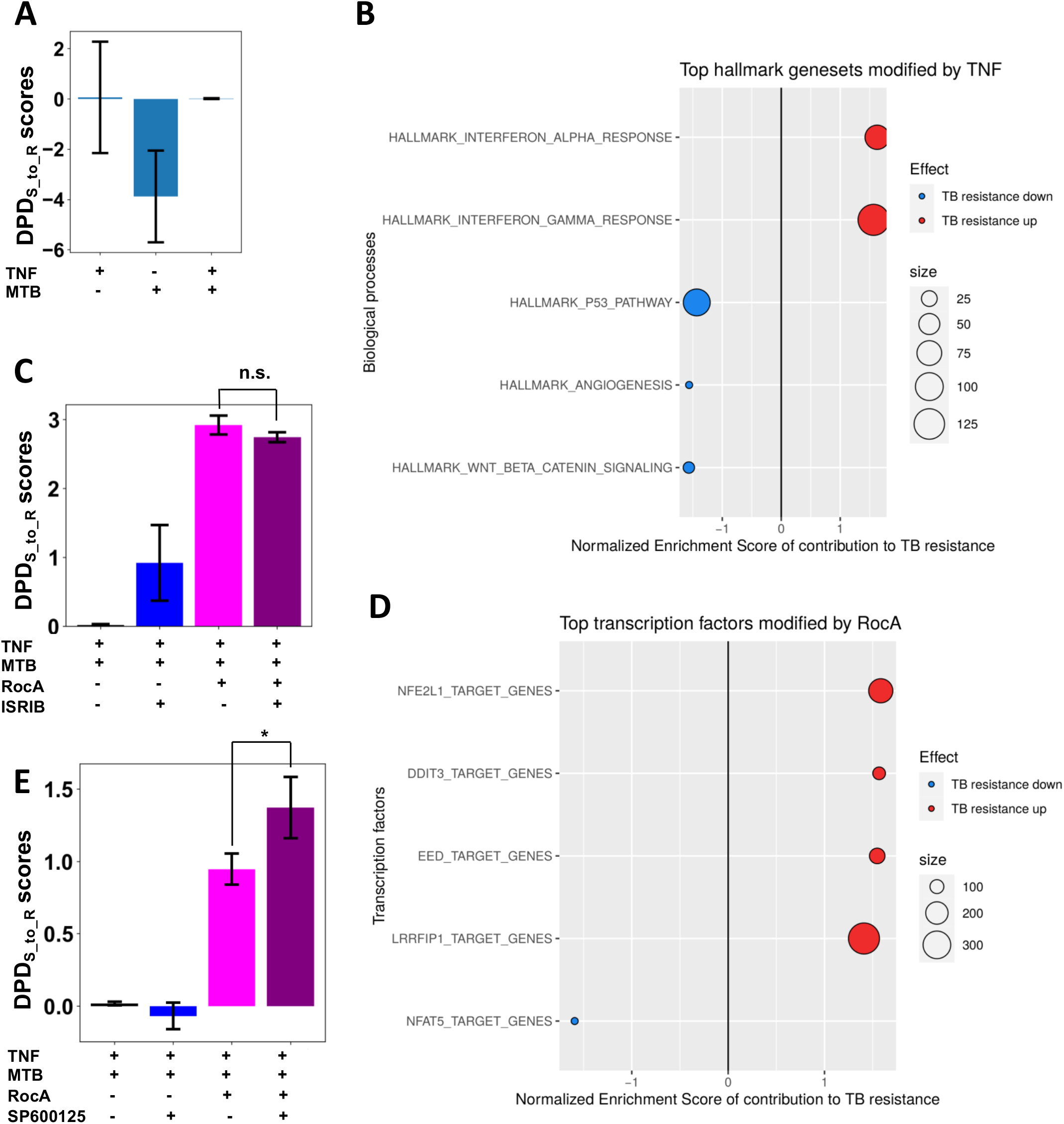
Effects of TNF and candidate compounds on the whole transcriptome responses of the susceptible macrophages to infection with virulent *Mycobacterium tuberculosis*. B6.Sst1S BMDMs were stimulated with 10 ng/ml TNF, infected with Mtb and, after phagocytosis, treated with candidate compounds for 3 days. The whole transcriptome analysis was performed using RNA-seq. **A**. The DPD_S_to_R_ scores for B6.Sst1S BMDMs naïve or stimulated with 10 ng TNF and infected with Mtb for 3 days. The separating hyperplane corresponds to the DPD_S_to_R_ = 0. **B**. Top GSEA Hallmark genesets driving DPD score in response to TNF. The projections of the log-fold gene expression changes into the STV_S_to_R_ have been used as the GSEA input for the GSEA hallmark gene set. Positive Normalized Enrichment Score (*red*) reflects improved Mtb resistance, negative Normalized Enrichment Score (*blue*) reflects impaired Mtb resistance. **C**. The DPD_S_to_R_ scores of B6.Sst1S BMDMs stimulated with 10 ng/ml of TNF and infected with Mtb (same as in A) and, after phagocytosis, treated with RocA, ISRIB and their combinations. The separating hyperplane corresponds to the DPD_S_to_R_ = 0. Positive DPD_S_to_R_ scores reflect the normalization of the B6.Sst1S BMDM transcriptome. **D**. Top GSEA predicted transcription factors driving DPD changes by RocA. The GTRD gene set (predicted transcription factor binding sites) was used for GSEA. Normalized Enrichment Score reflects improved (*red*) or impaired (*blue*) Mtb resistance. **E**. The DPD_S_to_R_ scores of B6.Sst1S BMDMs stimulated with 10 ng of TNF, infected with Mtb and treated with RocA, JNK inhibitor SP600125, or their combination. The test compounds were added to the infected BMDMs after phagocytosis of Mtb. The separating hyperplane corresponds to the DPD_S_to_R_ = 0 and positive DPD_S_to_R_ scores demonstrate the normalization of the infected B6.Sst1S BMDM transcriptome.

The RocA treatment led to dramatic changes of the entire transcriptome of TNF-stimulated and MTB-infected MFs, unlike ISRIB and JNKi, effect of which is much more targeted (**Supplementary Fig. 2**). RocA pushed S macrophage state across the separating hyperplane (making the DPD_S_to_R_ score positive, **Fig. 3C**). Importantly, RocA upregulated transcriptional targets of NFE2L1/2, the antioxidant defense transcription factors (**Fig. 3D**). Previously, we found that reactive oxygen species drive the aberrant TNF response in *sst1*-susceptible macrophages^32^. Thus, the induction of antioxidant defense may account for the rocaglate positive effect on TB resistance in our model.

Treatment with the inhibitor of the integrated stress response (ISR) ISRIB^42^ modulated the macrophage inflammatory and stress responses: the integrated stress response (ISR) genes, such as *Chac1*, were downregulated by ISRIB, whereas the interleukin 1 and interferon pathways were upregulated. The transcriptional responses of B6.Sst1S macrophages to the combination of rocaglate and ISRIB were similar to rocaglate alone and did not shift the DPD_S_to_R_ score closer to the resistant state (**Fig. 3C**). Perhaps, the effect of ISRIB becomes negligible in combination with RocA, because rocaglates reduce protein translation and prevent the proteotoxic stress^43^, thus, circumventing the eIF2*α*-mediated ISR.

In contrast, when we compared effects of RocA, a JNK inhibitor SP600125 and their combination, the DPD_S_to_R_ scores demonstrated a substantial synergy between RocA and JNKi (**Fig. 3E**). Although in this combination the signaling pattern was mainly determined by the rocaglate, yet the JNKi co-treatment further increased the DPD_S_to_R_ score. The analysis of the JNKi-specific contribution to the DPD_S_to_R_ score demonstrated the enhancement of the IL-1*β*, p38 signaling and the oxidative stress response pathways (**Supplementary Fig. 3**). Thus, RocA and JNKi boost two essential mechanisms of host resistance to Mtb – (i) macrophage responses to IL-1 and (ii) the antioxidant defense in a synergistic manner. We hypothesized that this synergy would allow using lower drug concentrations, thus avoiding potential toxicity, and improving their therapeutic effects.

### V. Testing cSTAR predictions in S macrophages infected with virulent Mycobacterium tuberculosis

To test combinatorial effects of compounds on Mtb-infected macrophages, we wanted to establish range of concentrations where individual compounds were biologically active and non-toxic during a 5 day Mtb infection assay. Therefore, we compared effects of RocA at 2.5 - 10 nM on Mtb-infected macrophage survival and Mtb loads during 5 days post infection. We observed no cell loss at 2.5 and 5 nM concentrations (**Fig.4A**). To monitor the bacterial loads, we used quantitative PCR based method for Mtb genome measurement^44^ and found that increasing RocA concentration did not improve the bacterial control (**Fig.4B**). To independently verify the effect of RocA at low concentration (3 nM) on Mtb control by S macrophages, we used acid fast fluorescent auramine-rhodamine staining to enumerate intracellular bacteria after membrane permeabilization. Of note, without permeabilization we detected no extracellular mycobacteria – a potential source of artifacts in the in vitro assays (**data not shown**). These data demonstrated that the RocA treatment significantly reduced the intracellular Mtb loads in TNF-stimulated B6.Sst1S macrophages and was not toxic during 5 days of the experiment (**Fig. 4C and D**). The treatment of BMDMs with RocA at 1 and 3 nM concentration showed dose dependent reduction in control of intracellular Mtb (**Supplementary Fig. 4)**. Therefore, to study potential synergy of RocA with ISRIB and JNK inhibitors, we used RocA at a suboptimal concentration (1nM).

**Figure 4.**
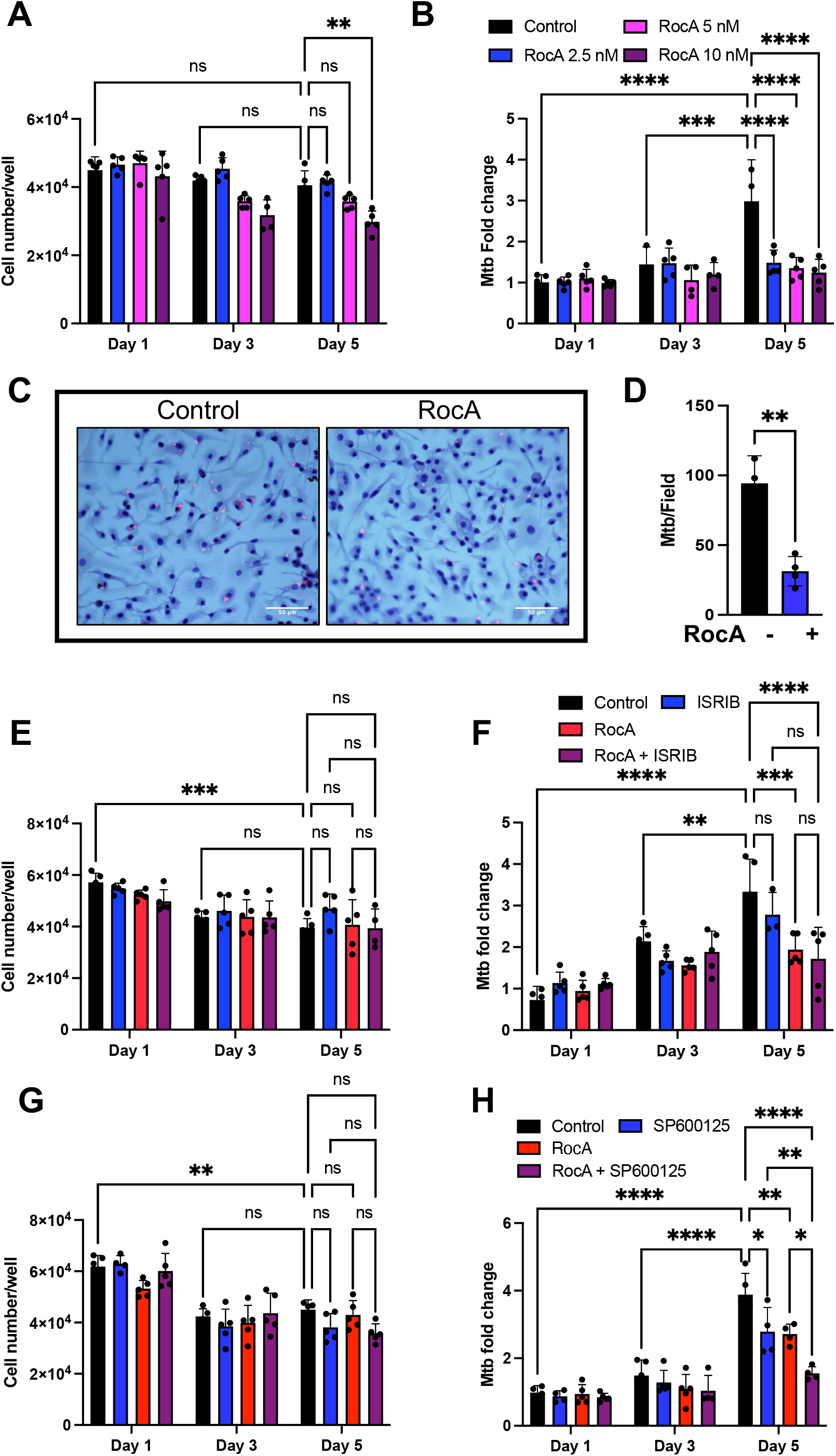
Effects of Rocaglates, ISRIB, and JNKi on BMDM survival and Mtb load. B6.Sst1S BMDMs were treated with 10 ng/ml TNF for 16 h and subsequently infected with Mtb at MOI=1 for 5 days. **A and B:** Cell survival (A) and Mtb load (B) after BMDMs were treated with different concentrations of RocA post phagocytosis. Cell numbers were determined using Celigo automated cytometer. Mtb loads were calculated as fold change by using quantitative genomic PCR (see Methods) at indicated time points. **C and D**. Intracellular MTB loads in cells treated with 3 nM RocA for 5 days. The cells were permeabilized with 0.05% tritonX-100 and Mtb stained using auramine-rhodamine. Scale bars=50 *μ*m. (D) Quantification of Mtb loads using ImageJ. **E and F**. Cell survival (E) and Mtb load (F) after BMDMs were treated with RocA (1 nM), ISRIB (10 *μ*M) or their combinations. Cell numbers were determined using Celigo automated cytometer. Mtb loads were calculated as fold change by using quantitative genomic PCR at indicated time points. **G and H**. Cell survival (G) and Mtb loads (H) after BMDMs were treated with RocA (1 nM), SP600125 (0.3 *μ*M) or their combination. Cell numbers were determined using Celigo automated cytometer. Mtb loads were calculated as fold change by using quantitative genomic PCR at indicated time points. The data represents the means ± SEM of four to five samples per experiment. The statistical significance was performed by two-way ANOVA using Bonferroni’s multiple comparison test (Panels A-B and E-H) and two-tailed unpaired t test (Panel C-D). Significant differences are indicated with asterisks (*, P < 0.05; **, P < 0.01; ***, P < 0.001; ****, P < 0.0001).

Treatment with ISRIB alone slightly improved the S macrophage survival, but did not improve their anti-mycobacterial activity (**Fig.4E and F**). Combining RocA with ISRIB also had no additional benefit in Mtb control, as compared to RocA alone (**Fig.4F)**, in agreement with the DPD score-based predictions **(Fig. 3B)**. In contrast, combination of RocA and JNK inhibitor SP600125 at suboptimal low concentrations (1 nM and 0.3 *μ*M, respectively) produced a cooperative effect without increased toxicity: five days post infection (p.i.), the Mtb loads in macrophages treated with individual compounds increased as compared to days 1 and 3 p.i., whereas the bacterial loads in macrophages treated with both compounds did not (**Fig. 4H**). (**Fig.4G**).

### VI. RocA increases antioxidant stress resilience of the susceptible macrophages

In-depth cSTAR analysis of transcriptomics data suggested that the primary mechanism by which RocA alone and in combination with JNKi improves Mtb resistance is by improving antioxidant properties of macrophages by upregulating genes of cells’ antioxidant defense. To test this suggestion, we analyzed biological markers of oxidative stress in macrophages treated with TNF, Mtb and drugs.

TNF stimulation and Mtb infection induces greater accumulation of lipid peroxidation (LPO) products, 4-HNE and MDA in B6.Sst1S BMDMs, as compared to B6 (Yabaji et al, in preparation). The RocA treatment reduced 4-HNE accumulation induced by TNF treatment (**Fig. 5A)** and Mtb infection (**Fig. 5B and C**). These data demonstrate that the rocaglate treatment corrected a defect in anti-oxidant response of B6.Sst1S macrophages.

**Figure 5:**
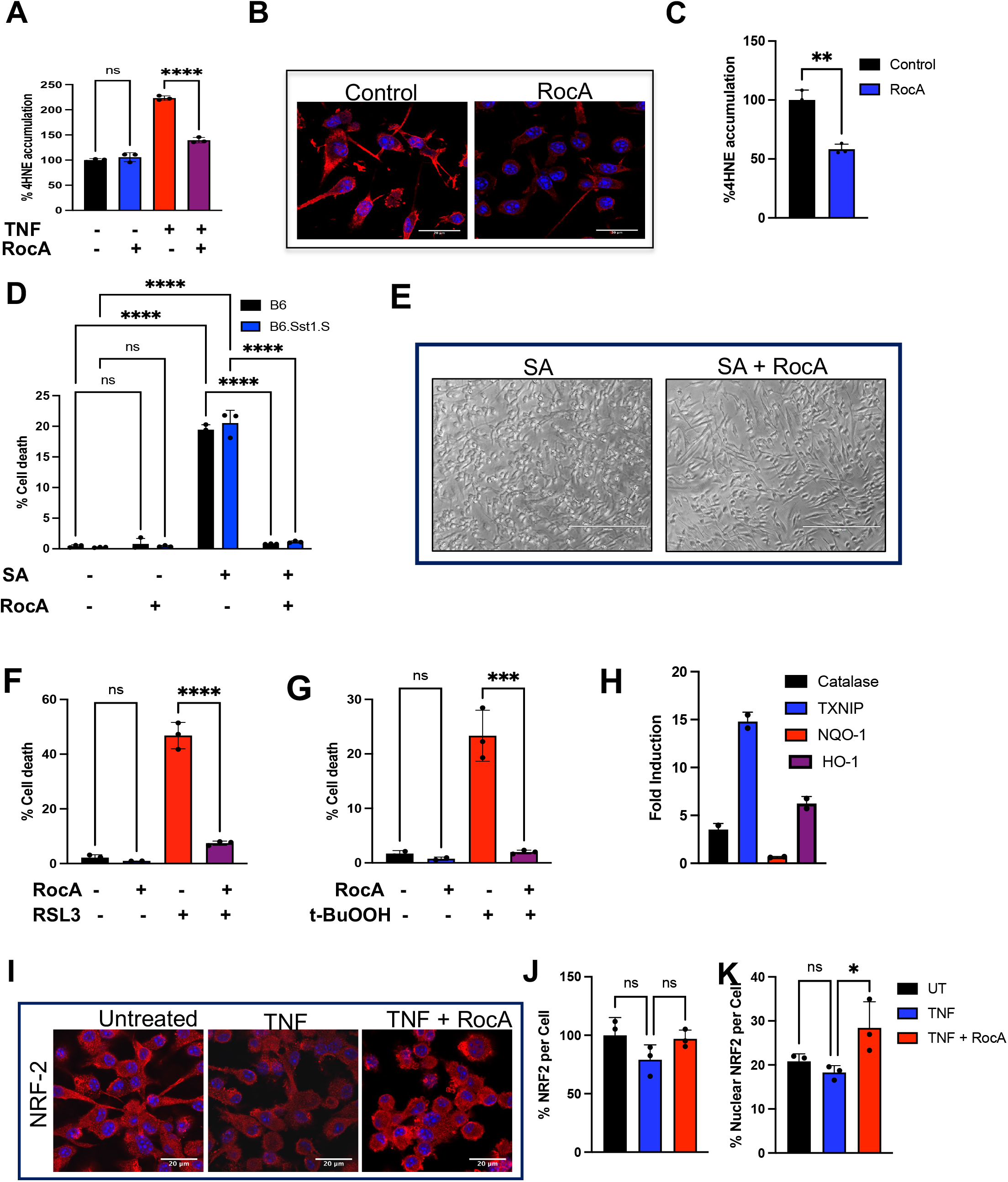
Rocaglate increases oxidative stress resilience of macrophages. **A**. RocA reduces the TNF-induced accumulation of lipid peroxidation product 4-HNE. B6.Sst1S BMDMs were treated with TNF alone or in combination with RocA (3 nM) for 48 h. The cells were stained using 4-HNE-specific antibody and accumulation of 4-HNE was measured using automated microscopy (Operetta CLS High Content Analysis System, PerkinElmer) and expressed as the % accumulation of 4-HNE, with TNF and RocA-untreated group as 100%. **B-C**. RocA reduces accumulation of 4-HNE during Mtb infection. B6.Sst1S BMDMs were pre-treated with TNF and subsequently infected with Mtb at MOI=1. The infected cells were treated with RocA (3 nM). Staining of 4-HNE with a specific antibody was assessed using confocal microscopy at day 5 post infection and expressed as the % accumulation of 4-HNE, with RocA-untreated group considered 100%. Scale bar – 20 *μ*m. **D-E**. RocA blocks cell death induced by sodium arsenate in both B6 and B6.Sst1S BMDM. (D) BMDMs from B6 and B6.Sst1S were pretreated with RocA (10 nM) for 4 h and subsequently cells were treated with 9 *μ*M sodium arsenate (SA) for 16 h. Cell death was detected by staining with Live-or-Dye™; 594/614 fixable viability dye (Biotium) and analyzed using Operetta CLS High Content Analysis System (PerkinElmer). (E) BMDMs from B6.Sst1.S were pretreated with RocA (3 nM) for 4 h and SA were added at 5 *μ*M concentration for 16 h. The brightfield images of macrophage monolayers were acquired at 20X magnification. Scale bar 200 *μ*m. **F**. RocA blocks cell death induced by GPX4 inhibitor. BMDMs from B6.Sst1S were pretreated with 30 nM RocA for 4 h. Inducer of ferroptosis, GPX4 inhibitor RSL-3 was added at 30 nM for 16 h. Percent cell death was determined as in D. **G**. RocA blocks cell death induced by lipid peroxidation. BMDMs from B6.Sst1S were pretreated with 10 nM RocA for 4 h; then 30 *μ*M t-BuOOH was added for 16 h. Percent cell death was determined as in D. **H**. Rocaglate induces expression of NRF2 target genes. B6.Sst1S BMDMs were treated with 100 nM CMLD010536 for 24 h. mRNA expression for catalase, *Txnip, Nqo1* and *Ho1* were measured using quantitative real-time PCR. **I-K**. RocA induces NRF2 nuclear translocation. B6.Sst1S BMDMs were treated with 10 ng/ml TNF alone or in combination with RocA (3 nM) for 30 h. The NRF2 protein level was determined using NRF2-specific antibody and confocal microscopy (I) the NRF2 total (J) and nuclear (K) levels were calculated using ImageJ. The NRF2 in untreated group was considered 100%. Scale bar 20 *μ*m. The data represent the means ± SEM of four to five samples per experiment. Statistical significance tests: (B, C) one-way ANOVA with Dunnett’s multiple comparison test, (E, J-K) one-way ANOVA with Bonferroni correction, (G) two-tailed unpaired t test, (H) two-way ANOVA with Bonferroni correction. Significant differences are indicated with asterisks (*, P < 0.05; **, P < 0.01; ***, P < 0.001; ****, P < 0.0001).

To study rocaglate effects on the macrophage oxidative stress resilience further, we treated macrophages with sodium arsenate (SA) – a potent inducer of oxidative stress. First, we found SA concentrations that induce chronic stress and kill macrophages over a period of 16-18 h in a concentration-dependent manner. The SA killing was not dependent on the macrophage *sst1* allele or TNF stimulation (**Supplementary Fig. 5A and B**). Pretreatment of macrophages with 3 nM RocA prevented the cell death induced by SA in both mouse strains (**Fig. 5D, E)**.

To test whether rocaglate can prevent an iron-dependent cell death, ferroptosis, we treated B6.Sst1S BMDMs with two ferroptosis inducers with different mechanisms of action: a small molecule inhibitor of Gpx4, RSL-3, and a lipid peroxidation inducer tert-Butyl hydroperoxide (t-BuOOH). At selected concentrations, both compounds induced macrophage death in TNF-independent manner. Pre-treatment with Roc A prevented ferroptosis in both cases (**Fig.5 F and G, Supplementary Fig. 5C**).

The treatment of macrophages with rocaglates for 24 h significantly upregulated the expression of antioxidant genes, which are known targets of NRF2: Catalase, TXNIP and heme oxygenase (HO-1) (**Fig. 5H**). Accordingly, we observed that the rocaglate pretreatment also increased nuclear translocation of NRF2 (**Fig. 5I and K)**, although the total cellular content of NRF2 was not affected **(Fig.5J)**. This observation is consistent with our previous finding based on proteomics analysis that rocaglate selectively downregulated KEAP1 protein – an E3 Ubiquitin ligase that targets NRF2 for degradation^29^and prevents its nuclear translocation. Thus, the rocaglate treatment broadly protects macrophages from oxidative damage by upregulating the anti-oxidant response.

## DISCUSSION

Although inflammation is a cornerstone of host resistance to infectious agents, during tuberculosis infection an exuberant inflammation is often associated with tissue damage and inefficient immunity. The diversity and connectivity of inflammatory pathways, however, provide an opportunity for selectively targeting pathological modules associated with inflammatory damage while preserving and restoring the protective immunity to Mtb.

This study outlines the design and implementation of a complete path from molecular characterization of macrophage transcriptional states to pharmaceutical interventions, an approach that is broadly applicable to diseases driven by pathological cell states. The key features of this strategy include (i) the use of primary cells from individuals representing resistant and susceptible phenotypes; (ii) focus on cell states rather than individual differentially expressed markers; (iii) analysis of multiple perturbations using cost-efficient targeted RNA-seq and (iv) use of cSTAR to quantify effects of individual perturbations that shifted the transcriptional states in the desired direction. In addition, this analysis enabled the imputation of synergistic drug effects that allowed us to reduce drug concentrations, focusing on desirable cell state transitions and mitigating the rocaglate toxicity at higher concentrations^29^. Ideally, this *in vitro* approach would predict bespoke interventions reducing pathological sequelae of dysregulated inflammation, while enhancing protective modalities.

To analyze the macrophage state transitions, we used a recently developed machine-learning approach cSTAR (cell State Transition Assessment and Regulation)^33^. This computational method uses multi-omics datasets to describe cell state transitions and build a cell State Transition Vector (STV) that indicates a path in a molecular dataspace to move a cell from one state to another state. Then the STV computes a cell Dynamic Phenotype Descriptor (DPD) that quantifies phenotypic changes in response to various perturbations. Using cSTAR, we identified small molecules that partially normalized the aberrant TNF response of *sst1*-susceptible macrophages, i.e., moved it closer to the pattern of relatively Mtb-resistant wild type macrophages, and calculated each gene’s contribution to the change in the DPD scores. These analyses also predicted synergistic effects of several candidate compounds.

We confirmed these predicted transcriptional states in the context of macrophage infection with virulent Mtb using whole transcriptome RNA-seq for in-depth analysis of cell states of macrophages treated with the selected candidate compounds and their combinations. The cSTAR analysis of the bulk RNA-seq data demonstrated that combination of RocA alone or in combinations with JNK inhibitor SP600125 improved the transcriptional response of the TNF-stimulated TB-susceptible macrophages to Mtb infection. This analysis also revealed that the rocaglate treatment boosted the antioxidant pathway gene expression. Importantly, the predicted transition between transcriptional states was also reflected in an improved control of the Mtb infection by the macrophages. We have experimentally validated the following: (i) the RocA treatment increased macrophage resilience to oxidative stress; and (ii) low concentrations of RocA and JNK inhibitor acted synergistically to improve control of virulent Mtb by the TB-susceptible macrophages. Our findings provide that targeting unresolving stress pathways associated with TB susceptibility is a promising strategy for host-directed TB therapies and provide candidates for the drug development.

## Materials and methods

### Experimental model and subject details

C57BL/6J mice were obtained from the Jackson Laboratory (ME, USA). The B6J.C3-Sst1C3HeB/FejKrmn (B6.Sst1S) mice were developed in our laboratory^45^ (available from MMRRC, stock # 043908-UNC). All experiments were performed with the full knowledge and approval of the animal ethics committee at Boston University (IACUC protocol number PROTO201800218).

### Reagents

Recombinant mouse TNF (Cat#315-01A) and IL-3 (Cat#213-13) were purchased from Peprotech. FBS (Cat# SH30396) for cell culture medium obtained from HyClone. Rocaglamide A (Cat#HY-19356), SP600125 (Cat#HY-12041) and RSL3 (Cat#HY-100218A) were purchased from MedChemExpress. ISRIB (Cat#SML0843) and tert-Butyl hydroperoxide (t-BuOOH, Cat# 458139) were obtained from Sigma-Aldrich. 4-HNE-(Cat#ab46545) and NRF2-(Cat#16396-1-AP) specific antibodies were purchased from Abcam and ProteinTech, respectively. All the primers and probes used were purchased from Integrated DNA technology (IDT). Middlebrook 7H9 and 7H10 mycobacterial growth media were purchased from BD and prepared according to manufactures instructions.

### Bacterial strains

The wild type *M. tuberculosis* H37Rv, Erdman (SSB-GFP, smyc’::mCherry)^46^ and *M. bovis* BCG Pasteur were grown at 37^°^C in Middlebrook 7H9 broth (BD Biosciences) or on 7H10 agar plates (BD Biosciences), respectively. Both solid and liquid media contained glycerol (0.5% v/v) and Tween 80 (0.05%). The MB7H9 broth was enriched using 10% ADC and MB7H10 agar was enriched using with 10% OADC. Mtb Erdman (SSB-GFP, smyc’::mCherry) was cultured in the presence of hygromycin B (50 *μ*g/ml).

### Primary Bone marrow derived macrophage (BMDM) cultures

Isolation of mouse bone marrow and BMDM growth and differentiation were performed as previously described^44^. For cell viability assays BMDMs were incubated with Live-or-Dye™; 594/614 Fixable Viability Stain (Biotium) at a 1:1000 dilution in PBS containing 1% FBS for 30 minutes at 37^°^C. After staining the samples were gently washed twice with 1% FBS in PBS and fixed with 4% Paraformaldehyde for 30 minutes. The fixed monolayers were washed, and the nuclei were stained using Hoechst 33342. The % of total and dead cells were determined by automated microscopy using Celigo microplate cytometer (Nexcelom) or Operetta CLS High Content Analysis System (PerkinElmer).

### Macrophage infection and quantification of intracellular bacterial loads

Single cell suspensions of BCG and Mtb were prepared as described in detail previously ^44^. BMDMs were infected with the mycobacteria in tissue culture treated 96 well plates at desired multiplicity of infection (MOI). The plates were centrifuged for 5 min at 200xg and incubated at 37^°^C with 5% CO_2_ for 1 h. Extracellular bacteria were inactivated by incubating the infected cell monolayers in medium containing 200 *μ*g/ml amikacin for 1 h, BMDMs were washed 3 times with 1% PBS containing 2% FBS and incubated at 37^°^C with 5% CO2. The intracellular bacterial loads were determined 1, 3 and 5 days post infection (p.i.) by a modified quantitative genomic PCR using Mtb and BCG specific primer/probes as described^44^.

### Imaging of mycobacteria in infected macrophages

The BMDMs were plated on cover slips placed in 24 well plates and infected with single cell suspensions of Mtb as above. The cells were fixed and permeabilized using 0.05 % Triton X-100 for 10 min, washed 3 times with 1X PBS for 5 min each. To visualize mycobacteria, the BMDM monolayers were stained using acid fast fluorescent staining with auramine O - rhodamine B at 37^°^C for 15 min, washed with 70% ethanol (3 times for 1 min each), counterstained with Mayer’s hematoxylin solution (Sigma-Aldrich Inc. St. Louis, MO, USA) for 5 min, and washed with distilled water. The bluing solution was added for 5 min to convert hematoxylin into an insoluble blue. Excess solution was washed with distilled water, then sections were dehydrated and mounted using Permount mounting medium.

For imaging the fluorescent reporter expressing Mtb Erdman (SSB-GFP, smyc’::mCherry), BMDMs were plated in 96 well glass bottom plate and infected with Erdman (SSB-GFP, smyc’::mCherry) at MOI=1. Cells were fixed and stained the nuclei with Hoechst 33342.

### Immunofluorescence microscopy

BMDMs were fixed with 4% paraformaldehyde for 10 min at room temperature and washed twice with 1X PBS. The permeabilization of cell was performed with 0.05% Triton X-100 and then blocked for 60 min with 1% BSA containing 22.52 mg/mL glycine in PBST (PBS+ 0.1% Tween 20). Cells were incubated with primary antibodies (NRF2-or 4-HNE-specific Ab) overnight at 4^°^C in 1% BSA, washed 3 times with 1X PBS for 5 min, and incubated with Alexa Fluor 594-conjugated Goat anti-Rabbit IgG (H+L) secondary Antibody (Invitrogen) in 1% BSA in dark for 1 h. The cells were mounted using ProlongTM Gold antifade reagent (Thermo Fisher Scientific) and Images were acquired using Leica SP5 confocal microscope. All images were processed using ImageJ software.

Alternatively, cells were plated in 96 well glass bottom plate (PerkinElmer) and processed and stained as above. After secondary antibody incubation cells were directly imaged and analyzed using Operetta CLS High Content Analysis System (PerkinElmer).

### RNA isolation and quantitative PCR

Total RNA was isolated using the RNeasy Plus mini kit (Qiagen). cDNA synthesis was performed using the SuperScript II (Invitrogen). Quantitative real-time RT-PCR (qRT-PCR) was performed with the GoTaq qPCR Mastermix (Promega) using the CFX-90 real-time PCR System (Bio-Rad). For calculating fold induction, the cycle threshold (Ct) of the test gene was normalized to the Ct of the internal control (18S) gene.

### Screen-seq methodology (adapted from Gupta, et al^39^)

Briefly, BMDM monolayers were prepared and treated in 96 well plates. Total RNA was extracted and from cell monolayers using a magnetic bead-based protocol using Sera-Mag SpeedBeads^™^ (GE Healthcare). Isolated RNA was DNase-treated with RQ1 DNase (Promega). The cDNA synthesis was performed within each well of a 96-well plate separately using EpiScript^™^ Reverse Transcriptase (EpiCentre Biotechnologies) using UMI-tagged oligo-dT primer with universal tail (**Suppl. Table 1**) using manufacturer’s instructions. Multiplex PCR was performed using Phusion U multiplex Master Mix (ThermoFisher, F562L). The multiplex products were used as templates to the second PCR using Phusion High-Fidelity DNA Polymerase (New England Biolabs Inc., M0530L) that barcodes each well/perturbation separately. These reactions use primers that target the universal tails (UT1 and UT2) of the readouts amplified in the first multiplex PCR and add a six-nucleotide barcode, a seven-nucleotide spacer and an Illumina primer dock. Combinatorial barcoding was achieved by using a pair of unique forward and reverse primers, which tag each sample with a unique barcode combination. These barcode combinations allowed pooling of samples in the subsequent steps of the assay. Once pooled, the readout library was cleaned up using magnetic beads at a bead to sample ratio of 1.2 to get rid of primer dimer bands <150bp in size. The sample was then carried over as a template into a third PCR reaction that added Illumina adapters.

### Screen-seq data analysis

After Screen-seq libraries were prepared as described above, they were sequenced at the UMass Boston and Center for Cancer Systems Biology (CCSB) sequencing core on Illumina HiSeq 2500 and MiSeq, respectively, using four-color reagent kits. From the P7 adapter end, 65nt were sequenced (Read 1), including one of the two barcodes for encoding plate wells and the UMI. From the P5 adapter the remaining 135nt were sequenced (Read 2), covering the second well-encoding barcode and the cDNA amplicon. Standard Illumina barcodes were used to distinguish individual plates within the overall pooled library, with demultiplexing before further processing. Reads were aligned using *bowtie2*^47^ against mm10 mouse genome assembly. The resulting BAM files were processed using custom Perl scripts to extract UMI-corrected counts for each gene and each well.

### Building the STVs and calculating the DPD scores

We first determined fold-changes in the targeted 46 RNA expression levels relative to the untreated control of TNF-treated B6 (R) and B6.Sst1S (S) macrophages. Each transcriptomics data point corresponds to a certain TNF dose for R or S macrophages, and these data can be perceived as points in the molecular data space of 46 dimensions of selected genes. In this transcriptomics dataspace, we used Support Vector Machines (SVM)^48^ to build two maximum margin hyperplanes that separate (i) the R and S macrophage states and (ii) transcriptomic patterns with no TNF and 50 ng TNF treatment. The SVM with a linear kernel from the scikit-learn python library^49^ was applied.

Next (part IV), we applied the SVM to distinguish transcriptomic patterns and build the separating hyperplanes in the bulk RNA space of 4305 dimensions. We used previously acquired transcriptomics data (accession number GSE115905) of responses to TNF R and S macrophages.

In this case, we have built two maximum margin hyperplanes (i) distinguishing R and S macrophage states, and (ii) distinguishing responses to stimulation by low (2 ng) and high (50 ng) TNF doses. The directions of changes in the transcriptomic patterns that correspond to different cell phenotypic features, (i) S and R macrophage TB susceptibility states and (ii) responses to TNF, are given by the two State Transition Vectors (STVs), the STV_S_to_R_ and STV_TNF_, respectively. Each STV is a unit vector defined as a vector of unit length that is normal to the corresponding separating hyperplane^33^.

After building the STVs, cSTAR calculates quantitative indicators of cell phenotypic states termed DPDs. The DPD scores describe the changes in the cell phenotypic features that would occur when the data point crosses on the separating hyperplanes. The absolute value of the DPD score is determined by the Euclidean distances from the data point to the separating hyperplane, while the sign of the DPD score indicates if the direction of the data point movement to cross the hyperplane is parallel or antiparallel to the corresponding STV^33^. The changes in the DPD scores following drug perturbations are readily calculated as the difference between the corresponding scores after and before the perturbation.

### Calculation of each gene contribution into the DPD scores and enrichment analyses

To determine the contribution of each gene into the changes in the DPD_S_to_R_ score following perturbations, we have calculated the projection of the log-fold change in this gene expression to the STV_S_to_R_. Mathematically, this is the dot product of the gene change and the STV, and their sum over all genes yields the change in the DPD score. Then, we used the top 400 contributing genes, which account for more than 80% of the DPD_S_to_R_ score change, to perform the enrichment analysis by the GeneOntology and STRING databases. We also input all gene names and their contributions into the DPD score into GSEA. Of note, the standard use of GSEA (where the GSEA input is the normalized read counts or log-fold-changes of differentially expressed genes) would not provide information about a transition from the S to R macrophage states. This information is contained in the STV_S_to_R_. Therefore, we used the projections of the corresponding gene expression changes into the STV_S_to_R_ as the GSEA input. Consequently, the normalized enrichment scores, which were generated by GSEA, now reflected how the changes in specific GSEA terms increased or decreased TB resistance of S macrophages.

### Quantification and statistical analysis

The Comparisons and statistical analysis were performed as indicated in each figure legend.

For calculating the mean values and errors of the DPD scores, outliers were removed from datasets based on analysis of correlation matrices and PCA plots of raw counts. Batch correction was performed for samples measured at different passages and days. Standard error of the mean for the DPD scores was estimated using raw counts of gene expression data. The Python code of building STVs, calculating the DPD scores for TNF treatments and drug perturbations, and estimation of errors of DPD scores is provided in the python worksheet ‘DPD_analysis.ipynb’.

To compare the multiple groups in the time course experiment or with two variables, a two-way analysis of variance (ANOVA) was used and corrected for multiple post hoc comparisons. The different comparisons used in our study including comparing all groups to each other, control to all the groups, or selected groups to each other. For comparisons of multiple groups with only one variable we used a one-way ANOVA and corrected for multiple post hoc comparisons. For comparisons of two groups, two-tailed paired or unpaired t tests were used. Except for sequencing data, statistical analyses were performed in the GraphPad Prism 9 software. We considered the *p* value ≤0.05 was statistically significant. *, p < 0.05; **, p < 0.01; ***, p < 0.001; ****, p < 0.0001.

## Acknowledgments

We thank Drs. Anwesha Nag and Kyomi Igarashi for their generous help in adapting Screen-seq protocol, Lorena Pantano for help with data analysis, Vadim Zhernovkov for useful discussions and assistance with data analysis. We thank Dr. Shumin Tan for advice and a generous gift of *M. tuberculosis* Erdman (SSB-GFP, smyc’::mCherry) replication reporter strain.

