## Supplementary Figures for "Cell state transition analysis identifies interventions that improve control of *M. tuberculosis* infection by susceptible macrophages"

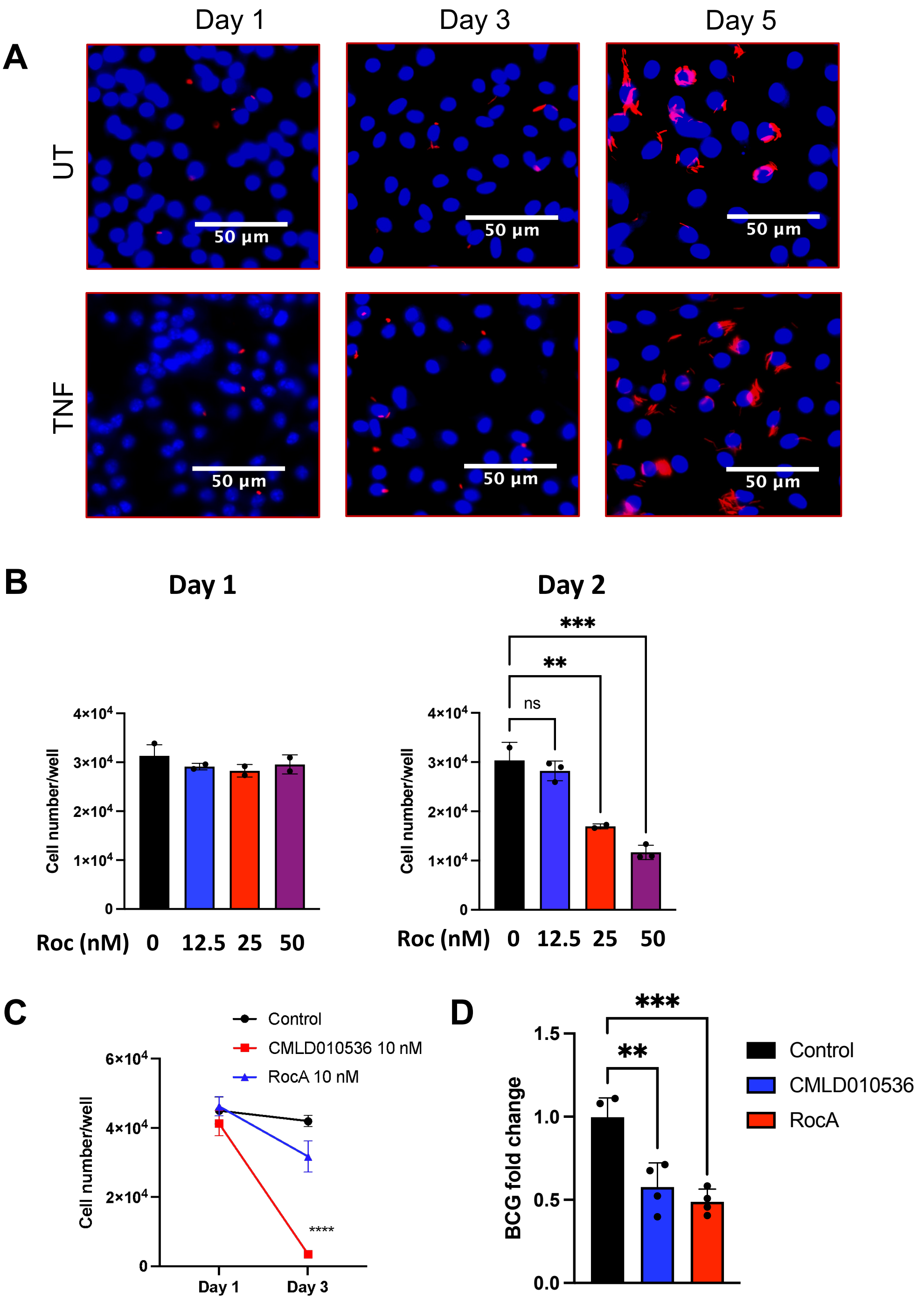


Supplementary Fig. 1

**Supplementary Fig. 1**

**A.** Fluorescent images of intracellular Erdman (SSB-GFP, smyc'::mCherry) (*red*) in B6.Sst1S BMDMs treated with TNF compared to untreated control. BMDMs were infected with Erdman (SSB-GFP, smyc'::mCherry) at MOI = 1 and imaged at 1, 3 and 5 days post infection. Nuclei (*blue*) – Hoechst 33342 counterstaining.

**B.** Increased toxicity of the synthetic rocaglate CMLD10536 for Mtb - infected B6.Sst1S BMDMs. Non-infected (left panel) and Mtb-infected (right panel), BMDMs were treated with increasing concentrations of CMLD010536 and cell numbers were analyzed using Celigo cytometer after Day 1 and Day 2 post treatment. At 25 – 50 nM CMLD10536 was not toxic for non-infected BMDMs, but its toxicity dramatically increased after infection with Mtb.

**C**. Plant-derived rocaglate Rocaglamide A (RocA) is significantly less toxic than CMLD010536. BMDMs were treated with TNF for 16 h followed by infection with Mtb at MOI=1 and the rocaglate treatment. The cell numbers were determined using Celigo automated cytometer at Day 1 and Day 3 post infection.

**D**. RocA and CMLD10536 demonstrated similar potency in enhancing macrophage control of *M.bovis* BCG. BMDMs were treated with TNF for 16 h and subsequently infected with BCG at MOI=1, and then treated with CMLD010536 or RocA at 3 nM concentration for 3 days. BCG loads were determined using quantitative genomic PCR and presented as fold change compared to Day 1. The p value $\leq 0.05$ was considered statistically significant.

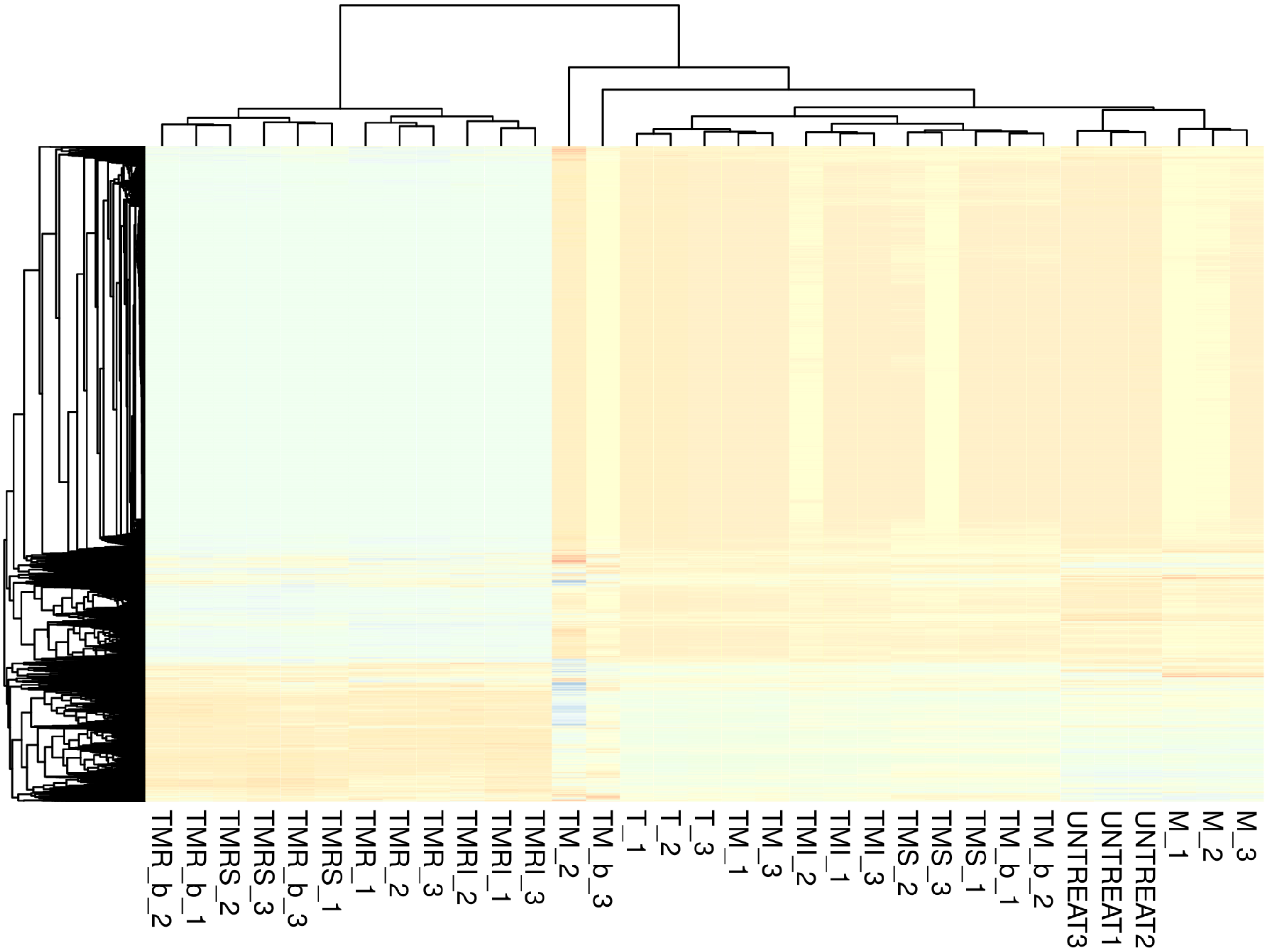


**Supplementary Fig. 2. Overview of RNA-seq perturbation data**

Heatmap of DESeq2 normalized counts of RNA-seq perturbation data. The sample name consists of the letters representing treatments applied, the number at the end of the sample name represents replicate number. “UNTREAT” represents no-treatment controls. “T” represents TNF treatment, “M” represents Mtb infection, “R” represents RocA treatment, “I” represents ISRIB treatment, “S” represents SP600125 treatment. Suffix “_b” represents second biological replicate of TM and TMR samples. Ward’s unsupervised clustering at the heatmap clearly shows that RocA treatment produces major effect on macrophage transcriptome.

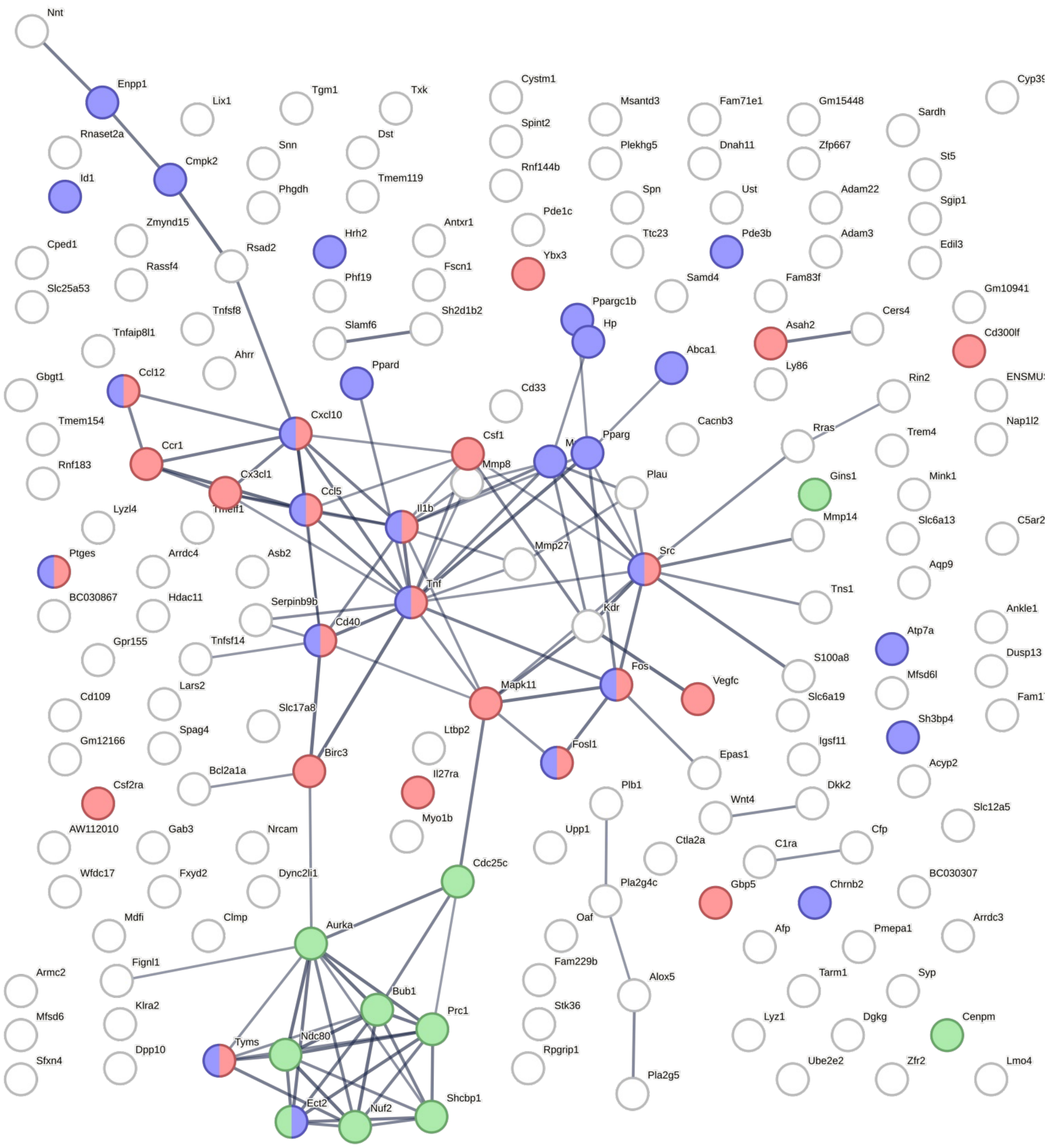


**Supplementary Fig. 3**

### Supplementary Fig. 3. Phenotypic effects of Rocaglates, JNK inhibitor and their combination

Genes that increase TB-resistance of Rocaglate-treated macrophages after addition of JNK inhibitor SP600125 were mapped to STRING database. Only high-confident interactions are shown (STRING score > 0.7). Red colour represents genes belonging to Gene Ontology term GO:0034097 “Response to cytokine”. Blue colour represents genes belonging to Gene Ontology term GO:1901700 “Response to oxygen-containing compound”. Green colour represents genes belonging to local network cluster identified by STRING as CL:2272 “Mixed, incl. condensed chromosome, centromeric region, and DNA replication”.

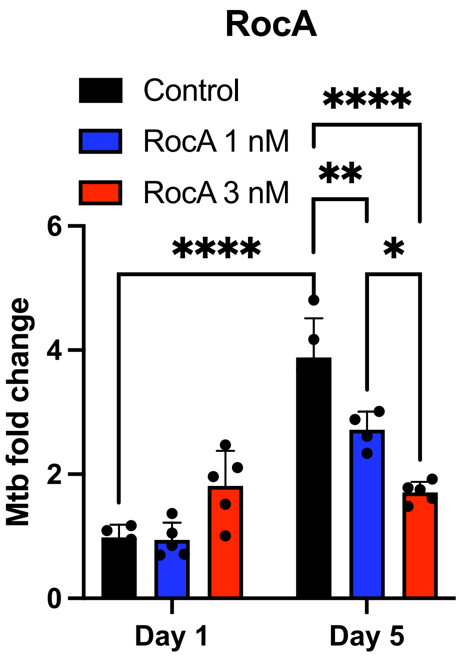


**Supplementary Fig. 4**

BMDMs were treated with TNF for 16 h followed by infection with Mtb at MOI=1. Post phagocytosis the cells were treated with RocA (1 and 3 nM). Mtb loads were calculated as fold change by using quantitative genomic PCR at indicated time points.
The p value ≤0.05 was considered statistically significant.

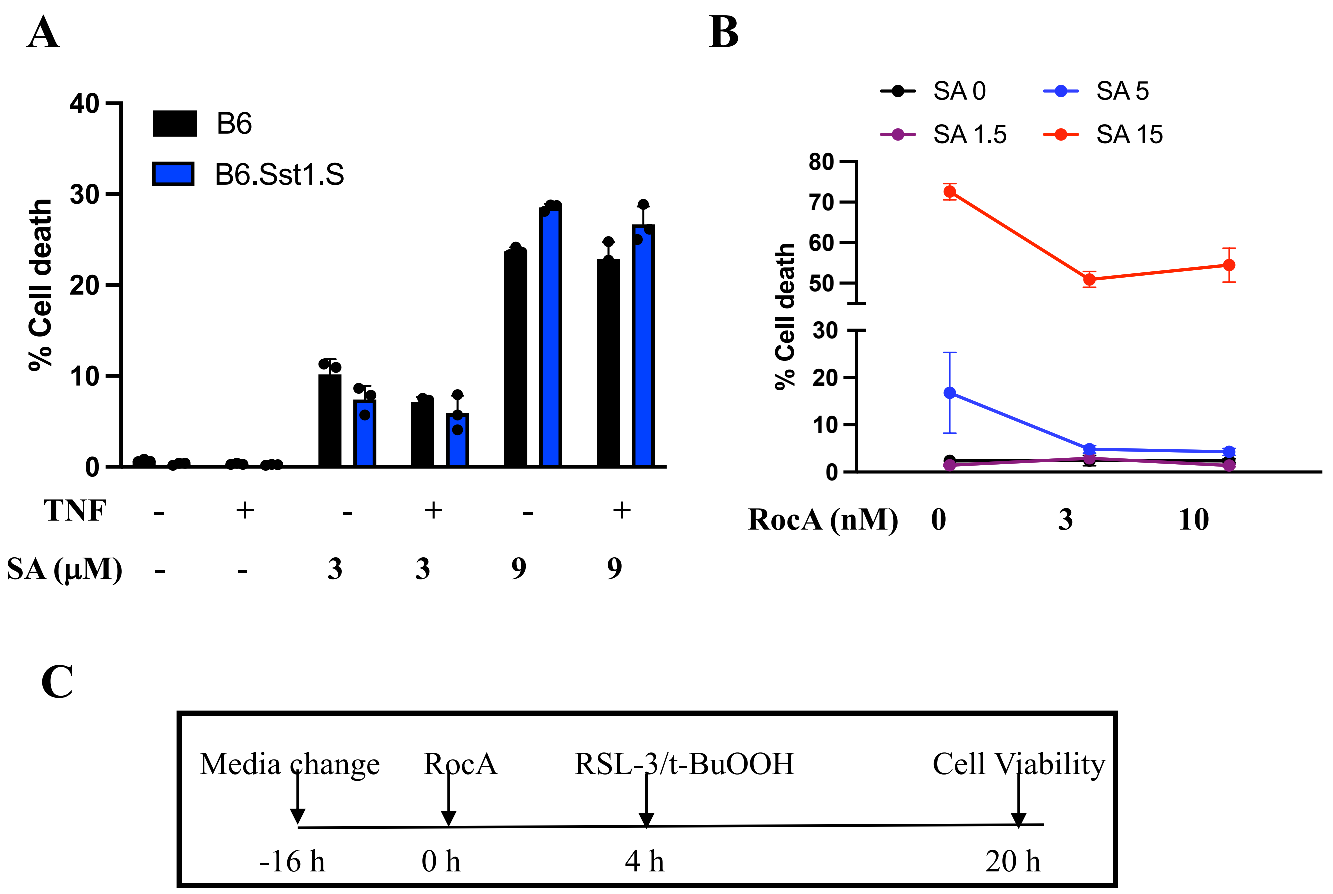


**Supplementary Fig. 5: Oxidative stress resilience by rocaglate.**

**A.** Sodium arsenate (SA) induces cell death in a concentration dependent manner in naïve and TNF-stimulated B6 and B6.Sst1S BMDMs. Cells were stimulated with TNF (10 ng/mL) for 2 h and incubated with SA at different concentrations for 16 h. Percent cell death was determined by staining cells with Live-or-Dye™ 594/614 fixable viability dye (Biotium) and analyzed using Operetta CLS High Content Analysis System (PerkinElmer). SA concentrations that induced rapid cell death were excluded.

**B.** RocA protects BMDM from oxidative stress. B6.Sst1.S BMDMs were pretreated with RocA at different concentration for 4 h and SA were added at 1.5, 5 and 15 µM concentration for 16 h. Percent cell death was determined as in A.
The p value ≤0.05 was considered statistically significant.

**C.** Experimental design for testing the RocA protection against ferroptosis inducers.
