## Supplementary Tables for "Cell state transition analysis identifies interventions that improve control of *M. tuberculosis* infection by susceptible macrophages"

**Supplementary Table 1**: **List of primers used for Screen-seq multiplex PCR**

| | |
|---|---|
| **UtailR_Oligo dT** | CGTGGAATCGCTAAAACGNNNNNNNNNNNTTTTTTTTTTTTTTTTTV |
| **Atf3** | CGTGGAATCGCTAATTGCAGAGCATTTCAGCTGGGAGA |
| **CD68** | CGTGGAATCGCTAATTGCCGCAGACGACAATCAACCTA |
| **Chac1** | CGTGGAATCGCTAATTGCGTCTGTCTGCTTGTTTGTCTGG |
| **G6pdx** | CGTGGAATCGCTAATTGCCCTCCTCACCAGATGGAAGA |
| **Itgb2** | CGTGGAATCGCTAATTGCCATAAAGGAAGACAGCAGTCTCAG |
| **Psmb10** | CGTGGAATCGCTAATTGCCAGAGAGCTGGCCGTTACC |
| **Rsad2** | CGTGGAATCGCTAATTGCAGCTCATGGACTGCCCTCT |
| **Sidt2** | CGTGGAATCGCTAATTGCCCCAGGAATGTGTTTCCTTC |
| **Rtp4** | CGTGGAATCGCTAATTGCTGTCTATTGCTGCATTTGCCCT |
| **Actb** | CGTGGAATCGCTAATTGCCCCACTCCTAAGAGGAGGATG |
| **Gsdmd** | CGTGGAATCGCTAATTGCTATGCTGAGGTGAAGGCTTGCT |
| **Klf4** | CGTGGAATCGCTAATTGCGCAAGCGCTACAATCATGGTCA |
| **Pcf11** | CGTGGAATCGCTAATTGCCAGCATCAAGATTTGCTGGCCT |
| **Six1** | CGTGGAATCGCTAATTGCAGTAGCCCAGCAGGAGTTTGAT |
| **Sp110** | CGTGGAATCGCTAATTGCACTCTCCAGCAGCAAATGACCT |
| **Tnfsf8** | CGTGGAATCGCTAATTGCTGGGCAAAGTGATCCTGTCTCT |
| **Apaf1** | CGTGGAATCGCTAATTGCCTGTAGCTGGCTAGTGAAGTG |
| **Atrnl1** | CGTGGAATCGCTAATTGCGCGGAGGTTGTTGGTACTT |
| **Bhlhe40** | CGTGGAATCGCTAATTGCGAGGCGAGGTTACAGTGTTTAT |
| **Cd33** | CGTGGAATCGCTAATTGCGTTGGGTCTAGCAATCTGTCTC |
| **Ch25h** | CGTGGAATCGCTAATTGCCCAGGACACAAGGTAGACATTC |
| **Chd7** | CGTGGAATCGCTAATTGCGCAATCCAGGAACGACTGATA |
| **Chn2** | CGTGGAATCGCTAATTGCCAAGGGTGAGGGTAGGAAAC |
| **Clk2** | CGTGGAATCGCTAATTGCGGGATATCAGTCGGTGACAATTA |
| **Ddit4** | CGTGGAATCGCTAATTGCCTGTTCGAGGCAGCTATCTTAC |
| **Dhcr24** | CGTGGAATCGCTAATTGCGGTCCCAGCCTTGTCTTATC |
| **Dusp10** | CGTGGAATCGCTAATTGCGAATCTGACGTATATGCCCTCATC |
| **E2f8** | CGTGGAATCGCTAATTGCCACACACACACACACACTCATA |
| **Fabp4** | CGTGGAATCGCTAATTGCACACTACAATAGTCAGTCGGATTT |
| **Fam49b** | CGTGGAATCGCTAATTGCGGAGTGAGTGAGGACTGATAAATG |
| **Fbxw7** | CGTGGAATCGCTAATTGCATTCACCAGGAGCCGTAACT |
| **Fzd1** | CGTGGAATCGCTAATTGCGTGTGCTCCTGTCTGGATTTAC |

| | |
|---|---|
| **Fzd7** | CGTGGAATCGCTAATTGCGATTGTCGGGAGGACAGATTAC |
| **Gbp4** | CGTGGAATCGCTAATTGCCTCGCAGATTTCTTGTCCTACT |
| **Golga2** | CGTGGAATCGCTAATTGCCCGACCTTAACCGTTCCAAT |
| **Hspa1b** | CGTGGAATCGCTAATTGCCCAGTAGCCTGGGAAGACATATAG |
| **Ifi205** | CGTGGAATCGCTAATTGCAGACCACAGTTTCGTCAAGG |
| **Ifrd1** | CGTGGAATCGCTAATTGCGCTATAGCGATCCTTCCAGTTT |
| **Igfbp4** | CGTGGAATCGCTAATTGCGGATTTGAACTCAGGACCTCTG |
| **Il7r** | CGTGGAATCGCTAATTGCGTGTCTGTCCGGTCATGTATTG |
| **Mmp13** | CGTGGAATCGCTAATTGCGACACAGCAAGCCAGAATAAAG |
| **Rhob** | CGTGGAATCGCTAATTGCCTGACCACACTTGTATGCTGTA |
| **Rnase4** | CGTGGAATCGCTAATTGCGAGAGATGCCTCTGTGGTTAAG |
| **Rnf213** | CGTGGAATCGCTAATTGCTGGTGCCCTAAAGTCCAATTC |
| **Serpine1** | CGTGGAATCGCTAATTGCGGCCACTCTGCATCTGTTATG |
| **Sh2d3c** | CGTGGAATCGCTAATTGCCTCTGAGCTCCATTCTTCCATC |
| **Smad3** | CGTGGAATCGCTAATTGCCCCAGTCCCTTCAACAGTATG |
| **Smad7** | CGTGGAATCGCTAATTGCGCTCGCTCGTATGATACTTTGA |
| **Stat1** | CGTGGAATCGCTAATTGCCAGGGCAAGACATCCACTTAC |
| **Wbscr27** | CGTGGAATCGCTAATTGCCAGTGCCAAACCTGGTAAGT |
| **Tnfrsf12a** | CGTGGAATCGCTAATTGCGGTGTCCAATTGCCCTATCT |

**Supplementary Table 2**: **List of compounds used and their concentration**

| Inhibitor (Pathway) | Name of inhibitor | Concentration used |
|---|---|---|
| p38 MAPK | SB203580 | 20 μM |
| JNK MAPK | SP600125 | 20 μM |
| PKR | C-16 | 2 μM |
| IFNAR1 | Anti-IFNAR Ab | 10 μg/mL |
| TBK1/ IKKε | BX795 | 2 μM |
| ROS | Butylated hydroxyanisole (BHA) | 100 μM |
| Integrated stress response | ISRIB | 10 μM |
| ATM | ATM inhibitor | 10 μM |
| ERK MAPK | U2016 | 20 μM |
| Myc | 10058-F4 | 50 μM |
| Brd4 | JQ-1 | 250 nM |
| PTEFb | Flavopiridol | 250 nM |
| TGFb | SB431542, SB525334 | 10 μM |
| Rocaglate | CMLD010536 | 30 and 100 nM |

**Supplementary Table 3: cSTAR analysis of targeted transcriptomics data- The DPD_TB values of WT and Sst1-mutant BMDMs treated with 10 ng TNF and different drugs for 24 hours.**

| **Condition** | **DPD_TB value** | **DPD_TB error** |
|---|---|---|
| Rocaglate (C10536) high dose | 11.214 | 0.596 |
| **Resistant macrophages** | **5.615** | **1.959** |
| Rocaglate (C10536) low dose | 2.544 | 1.659 |
| JNK inhibitor | -0.177 | 0.693 |
| TBK1 inhibitor | -1.686 | 0.498 |
| p38 and Myc inhibitors | -1.740 | 0.766 |
| ISRIB | -1.769 | 0.619 |
| IFNAR1 inhibitor | -1.827 | 0.469 |
| BHA antioxidant | -2.064 | 0.526 |
| p38 inhibitor | -2.338 | 0.671 |
| Myc and IFNAR1 inhibitors | -2.434 | 0.293 |
| PTEFb inhibitor | -2.672 | 0.539 |
| ERK inhibitor | -2.858 | 0.354 |
| p38 and IFNAR1 inhibitors | -2.864 | 0.122 |
| TGFb inhibitor | -3.195 | 1.153 |
| PKR inhibitor | -3.197 | 0.471 |
| Myc inhibitor | -3.589 | 0.452 |
| JQ1 (BET bromodomain inhibitor) | -3.972 | 0.804 |
| ATM inhibitor | -4.318 | 0.894 |
| **Sensitive macrophages** | **-6.630** | **0.188** |

**Supplementary Table 4: cSTAR analysis of targeted transcriptomics data- Top contributing genes to the STV_TNF and the STV_TB.**

| | |
|---|---|
| Genes that must be activated to switch to resistant phenotype (the highest positive STV_TB contributors/highest ranks) | E2f8, Gsdmd, Ifrd1, Stat1, Igfbp4, Fzd1, Ifi205 |
| Genes that must be suppressed to switch to resistant phenotype (highest negative STV_TB contributors/highest ranks) | Ch25h, Sh2d3c, Smad3, Fbxw7, Atrnl1, Itgb2, Il7r, Apaf1, Hspa1b, Fzd7 |
| Genes that are activated by TNF (highest positive STV_TNF contributors) | Serpine1, Ch25h, Gbp4, Fzd7, Hspa1b, Gsdmd, Fzd1, Rsad2 |
| Genes that are suppressed by TNF (highest negative STV_TNF contributors) | Tnfsf8, Igfbp4, Rnase4, E2f8, Wbscr27, Chn2, Smad3 |
